# Two gustatory receptors involved in host plant recognition of silkworm larvae

**DOI:** 10.1101/2022.09.12.507514

**Authors:** Haruka Endo, Kana Tsuneto, Takayuki Yamagishi, Dingze Mang, Katsuhiko Ito, Shinji Nagata, Ryoichi Sato

**Affiliations:** Department of Integrated Bioscience, Graduate School of Frontier Sciences, The University of Tokyo, Kashiwa, Chiba 277-8562, Japan; Graduate School of Bio-Applications and Systems Engineering, Tokyo University of Agriculture and Technology, Koganei, Tokyo 184-8588, Japan; Department of Science of Biological Production, Graduate School of Agriculture, Tokyo University of Agriculture and Technology, Fuchu,Tokyo 183-8509, Japan

**Keywords:** herbivorous insects, host plant selection, plant secondary metabolites, silkworm, gustatory receptors

## Abstract

Herbivorous insects can identify their host plants by sensing plant secondary metabolites as chemical cues. We previously reported the two-factor host acceptance system of the silkworm *Bombyx mori* larvae. The chemosensory neurons in the maxillary palp (MP) of the larvae detect mulberry secondary metabolites, chlorogenic acid (CGA), and isoquercitrin (ISQ), with ultrahigh sensitivity, for host plant recognition and feeding initiation. Nevertheless, the molecular basis for the ultrasensitive sensing of these compounds remains unknown. In this study, we demonstrated that two gustatory receptors (Grs), BmGr6 and BmGr9, are responsible for sensing the mulberry compounds with attomolar sensitivity for host plant recognition by silkworm larvae. Calcium imaging assay using cultured cells expressing the silkworm putative sugar receptors (BmGr4-10) revealed that BmGr6 and BmGr9 serve as receptors for CGA and ISQ with attomolar sensitivity in human embryonic kidney 293T cells. CRISPR/Cas9-mediated knockout (KO) of BmGr6 and BmGr9 resulted in a low probability of making a test bite of the mulberry leaves, suggesting that they lost the ability to recognize host leaves specifically. Electrophysiological recordings showed that the loss of host recognition ability in the Gr-KO strains was due to a drastic decrease in MP sensitivity toward ISQ in BmGr6-KO larvae and toward CGA and ISQ in BmGr9-KO larvae. Our findings unraveled that the two Grs, which have been regarded as sugar receptors, are molecules responsible for detecting plant phenolics in host plant recognition.

## Introduction

Many herbivorous insects have specialized on a limited range of plants as their hosts through ecological and evolutionary interactions with plants. Plants have various physical and chemical defense mechanisms to combat insect herbivory (War et al., 2012). For instance, plants have evolved a variety of secondary metabolites, most of which act as feeding deterrents and are often toxic to herbivorous insects (Fraenkel, 1959). In turn, herbivorous insects have adapted to some of the chemicals by developing chemosensory systems, enzymatic detoxication, and other mechanisms, which enabled them to consume a specific limited range of plants (Heidel-Fischer and Vogel, 2015; Simon et al., 2015). In parallel, they were able to identify host plants using plant secondary metabolites as sensory cues (Chapman, 2003).

Sensory exploration at the leaf surface is essential for host plant selection in larval feeding and adult ovipositioning (Chapman and Bernays, 1989; Renwick, 1989). We previously reported that the silkworm *Bombyx mori* larvae, well known for their preference for mulberry leaves, feed on the leaves utilizing a two-factor authentication system controlled by two taste organs and a set of chemoreceptors with differing sensitivities, operating sequentially in time (Tsuneto et al., 2020). The first step involves probing the leaf surface with the maxillary palp (MP), which contains ultrasensitive chemosensory neurons sensitive to a set of crucial mulberry compounds, including secondary metabolites (chlorogenic acid [CGA] and quercetin glycosides) and a phytosterol (β-sitosterol). The ultrahigh sensitivities in the attomolar and femtomolar ranges enable the larvae to detect trace amounts of these compounds at the dry leaf surface (Chapman and Bernays, 1989; Cheng et al., 2017) and identify the host leaves. If these three compounds are detected simultaneously, the larva makes a test bite, which leads to the second step. Another gustatory organ, the maxillary galea (MG), is used to probe the sugar content released by the test bite. Sucrose mainly induces continued biting, resulting in feeding. The two-factor host acceptance system is expected to be conserved among herbivorous insects because of their typical stepwise feeding behavior, composed of foraging, palpation at the leaf edge, test and persistent biting, and swallowing (Chun, 1972; Devitt and Smith, 1985). Although understanding the behavioral and neural basis of host acceptance in silkworm larvae has progressed, the molecular basis underlying MP’s ultrasensitive detection of the mulberry compounds remains unknown.

While, to our knowledge, no previous studies have identified lepidopteran chemosensory receptors responsible for host plant selection in larval feeding, two examples have been reported in adult oviposition. A gustatory receptor (Gr) PxutGr1 from the swallowtail butterfly *Papilio xuthus* is responsible for the sensitivity of tarsal taste sensilla to synephrine and the oviposition behavior induced by a mixture of synephrine and *chiro*-inositol (Ozaki et al., 2011). The two olfactory receptors (Ors) in *Plutella xylostella*, Or35 and Or49, are necessary and sufficient for detecting isothiocyanates and isothiocyanates-mediated oviposition preference for its host cruciferous plants (Liu et al., 2020). Meanwhile, some Grs have been reported as receptors for plant secondary metabolites in lepidopteran insects, although the importance in the ecology of the insect species is unclear. PrapGr28 from the cabbage butterfly, *Pieris rapae*, contributes to sinigrin sensitivity in adult tarsal taste sensilla (Yang et al., 2021), although sinigrin is a feeding stimulant for their larvae (Renwick and Lopez, 1999). PxylGr34 from the diamondback moth, *Plutella xylostella*, is involved in response to canonical plant hormones, brassinolide, and 24-epibrassinolide, a feeding and oviposition deterrent, in MG of larvae (Yang et al., 2020). In addition, BmGr16, BmGr18, and BmGr53 from *B. mori* respond to coumarin and caffeine in human embryonic kidney 293T (HEK293T) cells (Kasubuchi et al., 2018), but their role *in vivo* is unknown. Thus, several examples of relatively low-sensitive chemoreceptors for plant secondary metabolites in the foreleg of adult and MG of larvae have been reported. However, ultrahigh-sensitive chemoreceptors in larval MP underlying the first step in the two-factor host acceptance system are unknown.

We addressed the molecular basis of host recognition in silkworm larvae in the present study. *In vitro* calcium imaging in the cells suggested that BmGr6 and BmGr9 are candidates that confer MP ultrahigh sensitivity for CGA and ISQ. CRISPR/Cas9-mediated KOs demonstrate that the two Grs are responsible for host recognition by sensing CGA and/or ISQ.

## Results

### BmGr6 and BmGr9 serve as ultrahigh-sensitive receptors for CGA and ISQ as well as sugar receptors in HEK293T cells

We started with phylogenetic analysis to find closely related silkworm Grs with PxutGr1 as candidate receptors for CGA and ISQ because PxutGr1 is the only lepidopteran Gr reported as a receptor for plant phenolics (Ozaki et al., 2011). Phylogenetic analysis of lepidopteran Grs indicates that PxutGr1 belongs to the same lineage as BmGr9 and BmGr10, known as fructose and *myo*-inositol receptors, respectively (Kikuta et al., 2016; Sato et al., 2011) (Fig. 1). Based on the phylogenetic analysis, we focused on BmGr9, BmGr10, and other five Grs (BmGr4-8) in the neighboring putative sugar receptor clade (Wanner and Robertson, 2008).

**Fig. 1.**
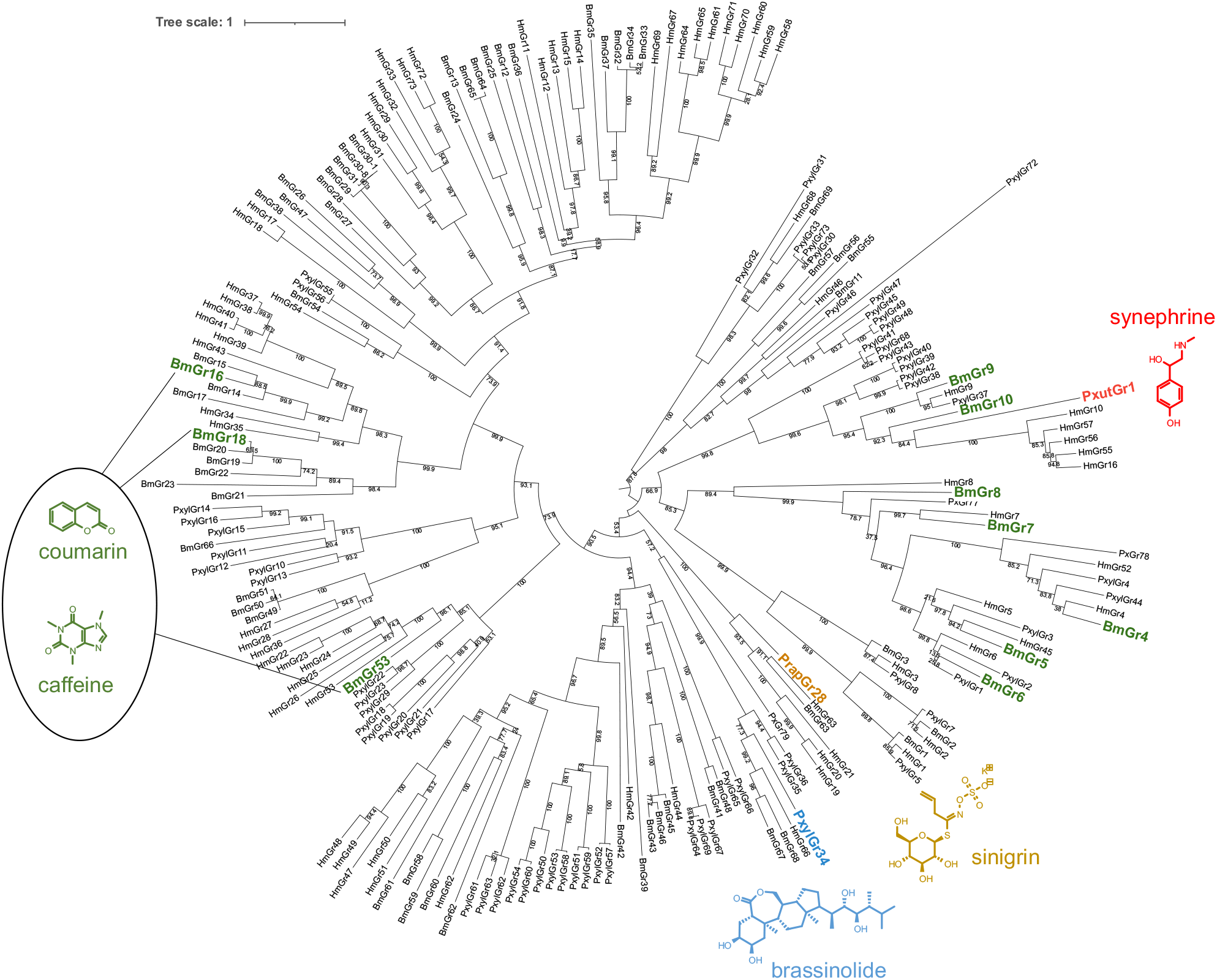
Phylogeny of the gustatory receptors in some lepidopteran insects. The combination of gustatory receptors and plant secondary metabolites as their ligands were highlighted with the same colors as reported in previous studies [12,14,16,17]. Bm: *B. mori* (Guo et al., 2017); Pxyl: *P. xylostella* (Engsontia et al., 2014); Hmel: *H. melpomene* (Briscoe et al., 2013).

To test whether these seven Grs can confer silkworm MP chemosensory neurons attomolar sensitivity to CGA and ISQ, we performed calcium imaging in HEK293T cells. Of the seven Grs-expressing cells, BmGr6-, BmGr8-, and BmGr9-expressing cells responded to 100 aM CGA, and BmGr6-and BmGr9-expressing cells responded to 100 aM ISQ (Fig. 2A-C). We recorded a dose-dependent response of BmGr6, BmGr8, BmGr9, and BmGr10 (Fig. 2D and E) and determined the EC_50_ values (Table S1). Although BmGr6-and BmGr9-expressing cells responded to <10 aM CGA and ISQ, BmGr8-expressing cells did not respond to 10 aM CGA (Fig. 2D). Based on these results, we focused on and further characterized BmGr6 and BmGr9 as candidates conferring MP neurons attomolar sensitivities toward CGA and ISQ. BmGr6-and BmGr9-expressing cells did not respond to β-sitosterol up to 2 µM (Fig. S1).

**Fig. 2.**
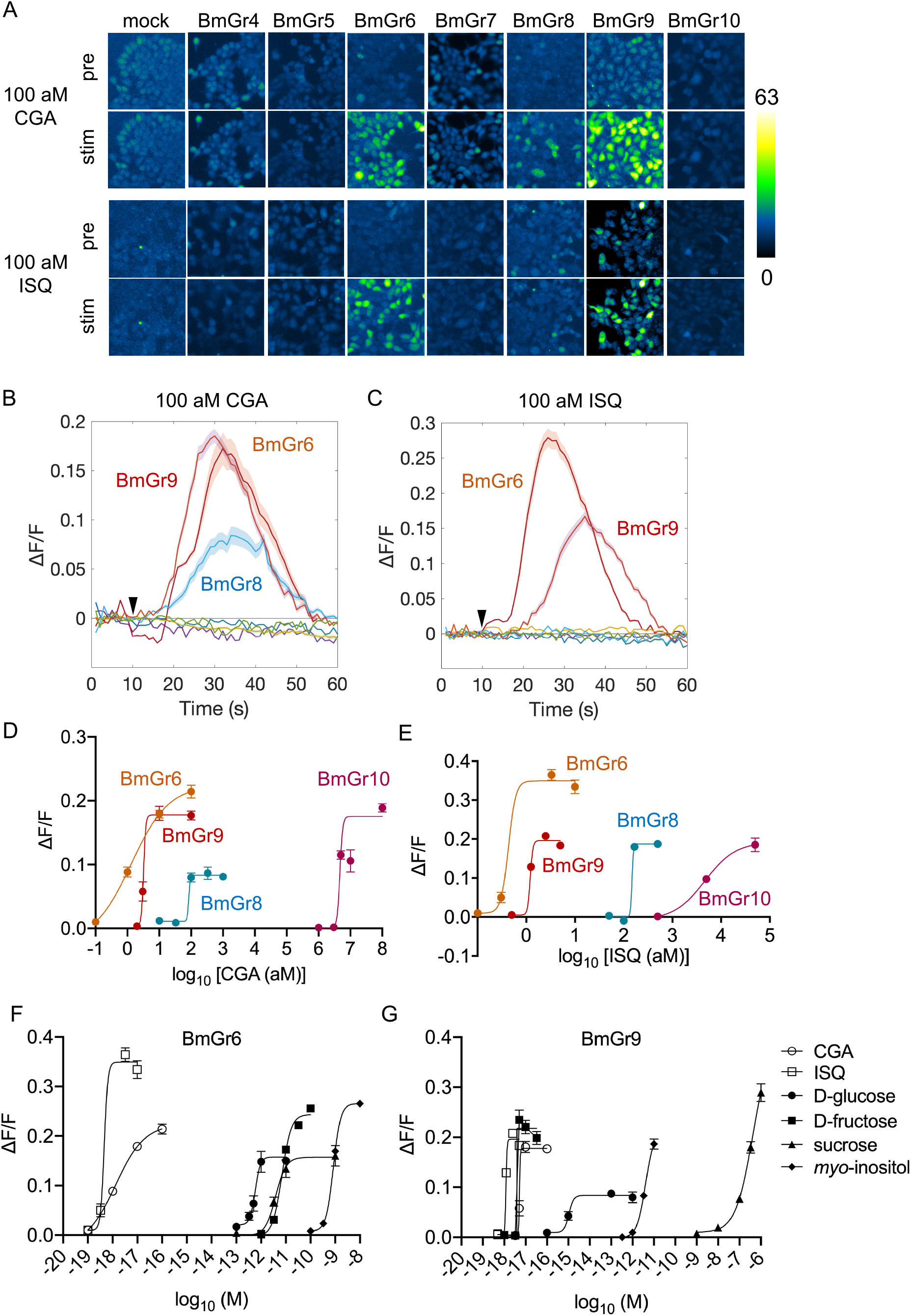
Calcium imaging of BmGrs-expressing HEK293T cells. (A) The response of BmGr-expressing cells to 100 aM chlorogenic acid (CGA) and isoquercitrin (ISQ). The representative cell images before (pre) and after (stim) stimulation. Fluorescent intensity is color-coded according to the scale bar. (B and C) Time course of responses (ΔF/F) toward 100 aM CGA (B) and ISQ (C) (n = 30). The arrowheads indicate the timing of the ligand application. (D and E) Dose-dependent responses of BmGr6, BmGr8, BmGr9, and BmGr10 toward CGA (D) and ISQ (E). (F and G) Dose-dependent responses of BmGr6 (F) and BmGr9 (G) toward CGA, ISQ, and sugars (B–G). Data are represented as the mean ± SEM.

Since BmGr6 is regarded as a putative sugar receptor (Wanner and Robertson, 2008) and BmGr9 was reported as a fructose receptor (Sato et al., 2011), we investigated their response to major sugars present in the mulberry leaves (D-fructose, D-glucose, sucrose, and *myo*-inositol) (Kikuta et al., 2016). Surprisingly, BmGr9-expressing cells responded to D-fructose with aM sensitivity (Fig. 2C and D), whereas they exhibited relatively lower sensitivities to D-glucose, sucrose, and *myo*-inositol (Fig. 2E and F). BmGr6-expressing cells showed pM to nM sensitivity to four sugars (Fig. 2C-F). These results indicate that the two Grs are highly tuned to CGA and ISQ, whereas BmGr9 also exhibits a high affinity with D-fructose.

### BmGr6 and BmGr9 are expressed in subsets of MP chemosensory neurons

According to the RNA-seq data from a previous study, BmGr6 and BmGr9 are highly expressed in the MP (Guo et al., 2017). To confirm this at the protein level, we performed immunohistochemistry using anti-BmGr6 and anti-BmGr9 antisera. Each sensillum in MP contains three to five chemosensory neurons whose somas are usually clustered in the second segment (Fig. 3A and 3B) (Asaoka, 2003). We observed three BmGr6-antiserum-positive MP neurons (Fig. 3C). In the two representative cryosections, two of the three neurons are close to each other (Fig. 3C), suggesting that the two neurons are contained in the same sensillum. We observed seven BmGr9-antiserum-positive MP neurons, three of which are close to each other (Fig. 3D). Thus, BmGr6 and BmGr9 are expressed in at least three and seven neurons in the MP of silkworm larvae.

**Fig. 3.**
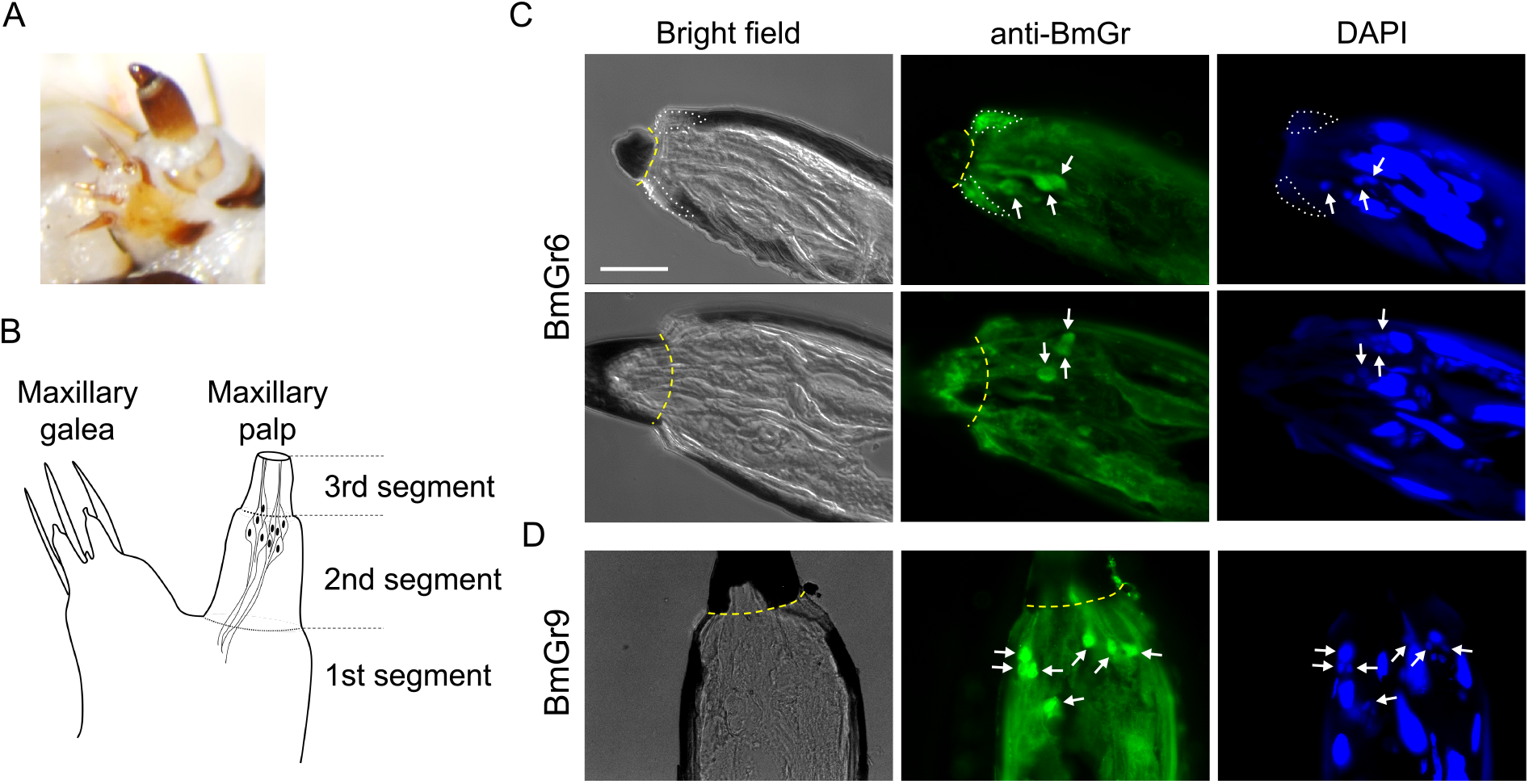
Immunostaining using maxillary palp (MP) BmGr6 and BmGr9 antibodies. (A and B) Image (A) and schematic diagram (B) of the maxilla of a silkworm larva. (C and D) Expression of BmGr6 (C) and BmGr9 (D). Arrows indicate somas of immunoreactive sensory neurons. The areas surrounded by broken white lines indicate the autofluorescence of the cuticle. Yellow broken lines indicate the border between MP’s second and third segments. Scale bar: 25 μm.

### BmGr6-KO and BmGr9-KO larvae lose their ability to recognize mulberry leaves specifically

To test whether the two Grs are involved in host recognition *in vivo*, we generated BmGr6-KO and BmGr9-KO silkworm strains via the CRISPR/Cas9 system. We designed single-guide RNAs (sgRNAs) targeting the exons 5 and 3 of the *BmGr6* and *BmGr9* genes (Fig. 4A). By injecting the sgRNAs and the Cas9 protein into eggs, we successfully established mutant strains carrying homozygous null mutations of *BmGr6* and *BmGr9* (material and methods for details). In both strains, premature stop codons were generated via indel mutations (Fig. 4A and S2).

**Fig. 4.**
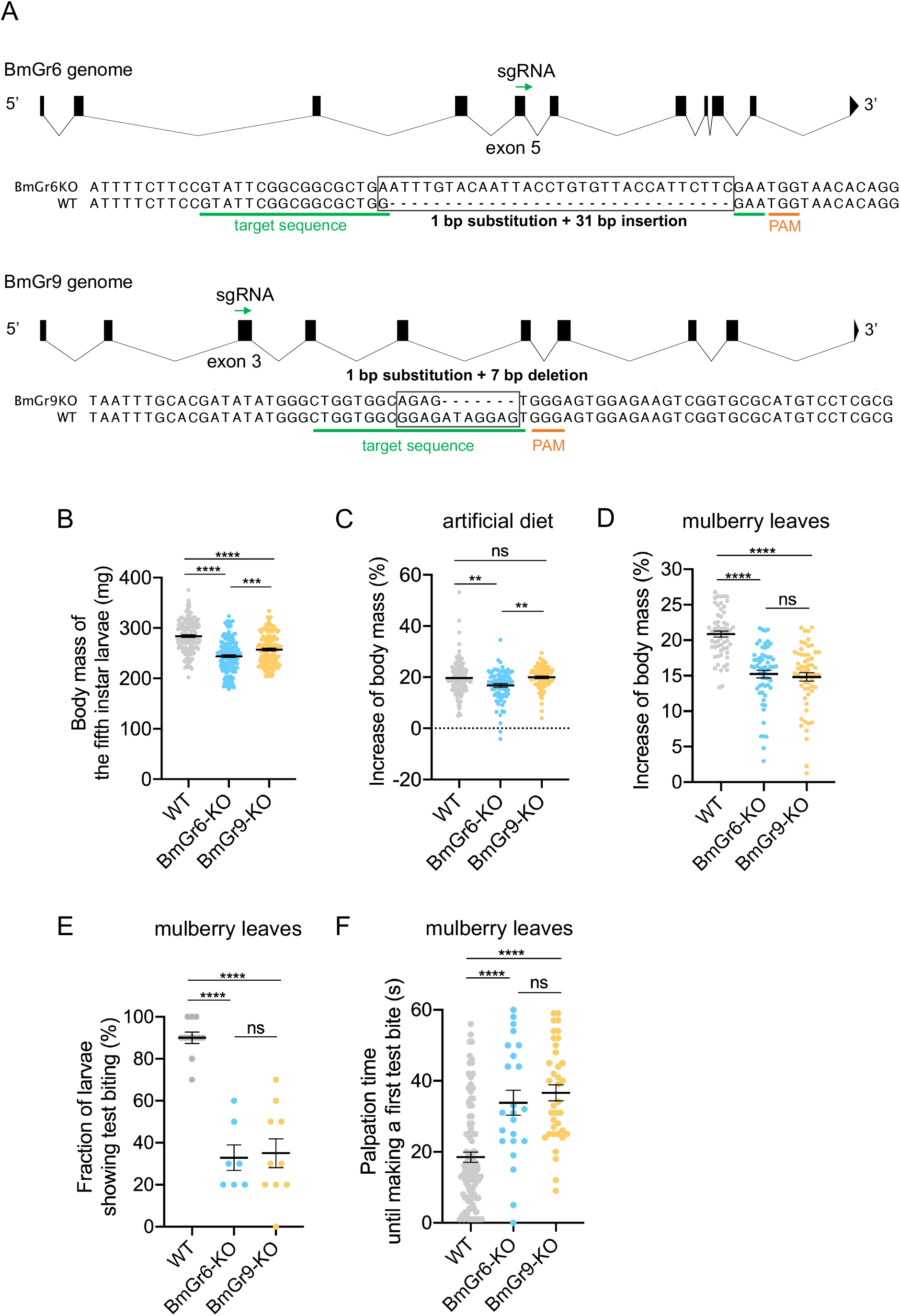
Generation and characterization of BmGr6-KO and BmGr9-KO larvae. (A) Gene structures, sgRNA target sites, including the adjacent protospacer motif (PAM), and obtained CRISPR/Cas9-mediated mutations of *BmGr6* and *BmGr9*. Gene structures were drawn using Exon-Intron Graphic Maker (http://wormweb.org/exonintron). (B) The body mass of freshly molted larvae (Day 0 of the fifth instar) (n = 140–175). (C and D) The increase in larval body mass after 1.5 h feeding of artificial diets (C) (n = 75–117) and mulberry leaves (D) (n = 60). (E) The proportion of larvae showing test biting toward mulberry leaves. The biting of ten larvae was observed for one minute. Experiments were repeated as independent biological replicates (n = 7–11). (F) Palpation time until the first bite toward mulberry leaves (n = 23–99). Data are represented as the mean ± SEM. Statistical analysis was performed by one-way ANOVA followed by the Tukey post hoc test. “ns” indicates no significant difference; an asterisk indicates a significant difference (\*\**P* < 0.01; \*\*\*\**P* < 0.0001).

The body mass of the fifth instar larvae of BmGr6-KO and BmGr9-KO strains, which have been fed with artificial diets since hatching, were ∼86% and ∼91% of the body mass of wild-type (WT) larvae (Fig. 4B). Since BmGr6 and BmGr9 are expressed in neurosecretory cells in the brain, enteroendocrine cells in the midgut, and other cells in a diversity of organs (Mang et al., 2016; D. Mang et al., 2016), many factors could influence the body mass in the mutant lines in addition to gustation. To directly see the effects of Gr knockouts on feeding, we conducted 1.5 h short-term feeding assay using artificial diets and mulberry leaves. When feeding on leaves, the larvae must recognize host leaves by MP to initiate feeding (the first step of the host acceptance system) (Tsuneto et al., 2020). This step can be skipped when feeding artificial diets with abundant sugars on their surface are available. The feeding amount of artificial diets by BmGr6-KO larvae was ∼85% of the WT larvae (Fig. 4C), suggesting that the low body mass of BmGr6-KO larvae is due to the low feeding amount of artificial diets. The feeding amount of BmGr9-KO larvae is comparable with that of WT (Fig. 4C). Metabolic factors and factors that affect long-term feeding, such as the interval of feeding, might cause the low body mass of BmGr9-KO larvae. The feeding amounts of mulberry leaves by the two mutant larvae were significantly lower than the WT (Fig. 4D). To directly observe the ability of host recognition, we next observed test bites toward mulberry leaves. Only 33% and 35% of BmGr6-KO and BmGr9-KO larvae made test bites within 1 min, whereas 90% of WT larvae made test bites (Fig. 4E). This low probability of test biting corresponds to those of some nonhost leaves in WT larvae (Tsuneto et al., 2020), suggesting that the two mutant strains lose their ability to recognize mulberry leaves. Even when BmGr6-KO and BmGr9-KO larvae were making test bites within 1 min, and their palpation time until making a first test bite was significantly longer than that of WT larvae (Fig. 4F), suggesting that even if mutant larvae made a test bite, they were not able to recognize mulberry leaves immediately. These results indicate that BmGr6 and BmGr9 are essential for host recognition *in vivo*.

### BmGr6 and BmGr9 are responsible for sensing ISQ and both CGA and ISQ, respectively: *in vivo*

We performed electrophysiological recordings from MP to test whether the two Grs are involved in CGA and ISQ sensing *in vivo*. The MP neurons of BmGr6-KO larvae started responding to CGA from 10 aM and showed almost the same sensitivity as WT larvae (Fig. 5A and B); they did not respond to ISQ up to 1 µM (Fig. 5A and C). This suggests that the ISQ sensitivity of BmGr6-KO larvae was >10^10^-fold lower than the MP neurons of WT larvae. By contrast, both CGA and ISQ sensitivities of BmGr9-KO larvae were ∼1,000-fold lower than those of WT larvae (Fig. 5A-C), suggesting that BmGr9 is required for attomolar sensitivities of both CGA and ISQ. We previously reported that the β-sitosterol sensitivity of MP neurons is at the femtomolar level (Tsuneto et al., 2020). The number of spikes toward 10 fM β-sitosterol was comparable between WT, BmGr6-KO, and BmGr9-KO strains (Fig. S3), suggesting that MP of the two mutant strains preserved the sensitivity to β-sitosterol. Note that although spikes with multiple shapes derived from different MP chemosensory neurons were expected, we did not observe such spikes but rather observed spikes that are similar to those of single neurons. This may be due to whole MP sensilla recording. Single MP sensillum recording and other imaging methods should be established to clarify how many chemosensory neurons respond. Taken together, BmGr6 and BmGr9 are required for the attomolar sensitivity to ISQ, and BmGr9 is required for attomolar sensitivity to CGA; nevertheless, BmGr6 is not involved in CGA sensing *in vivo*. Unlike *in vitro* study observations in Fig. 2, these results indicate different roles of BmGr6 and BmGr9 *in vivo*.

**Fig. 5.**
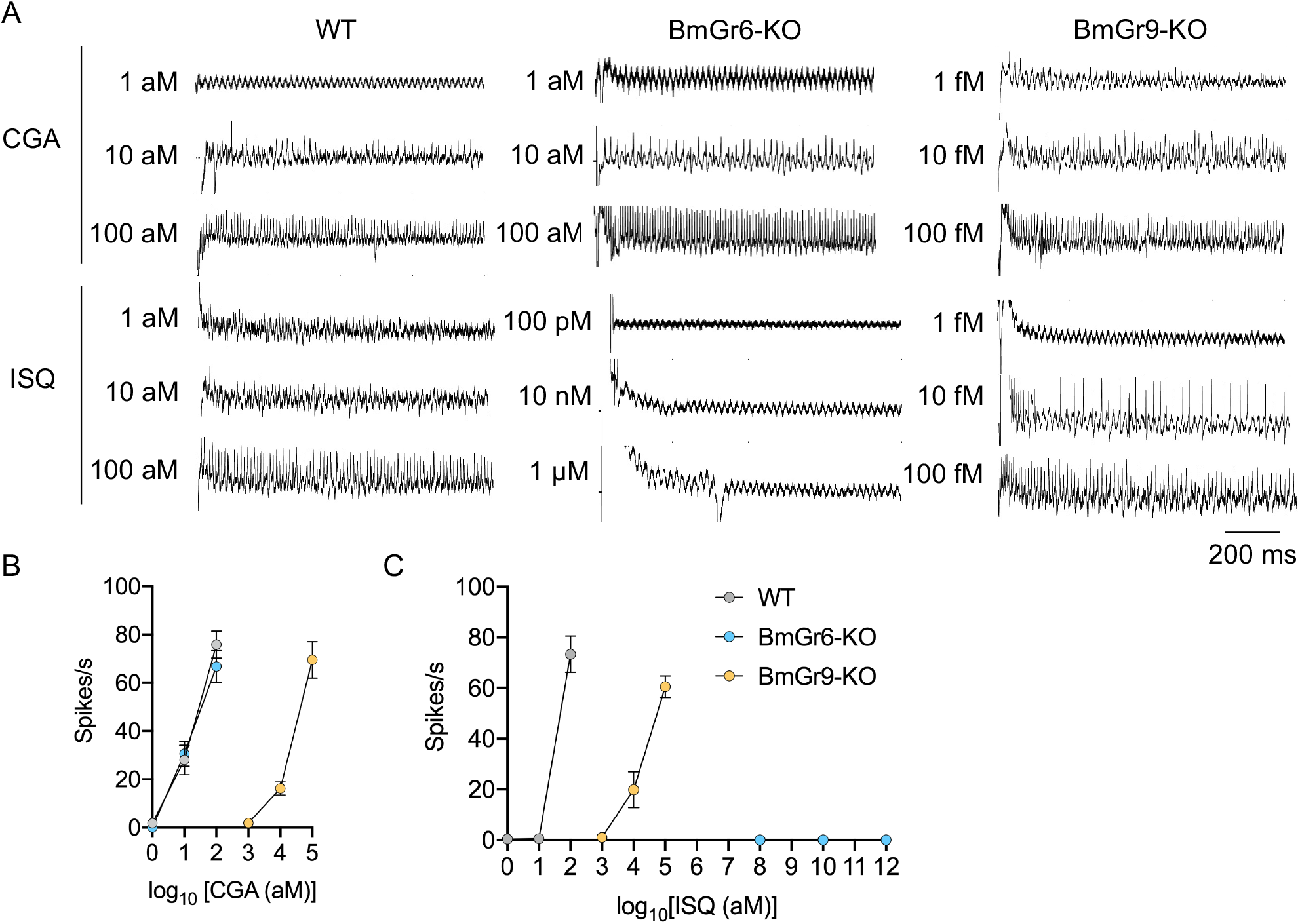
Electrophysiological recordings of the maxillary palp (MP) in BmGr6-KO and BmGr9-KO larvae. The representative traces (A) and frequencies of action potentials from the whole MP sensilla in response to chlorogenic acid (CGA) (B) and isoquercitrin (ISQ) (C). Spikes were counted from 0.1 to 1.1 s after the onset of stimulation. Error bars are represented as SEM (n = 6–8).

## Discussion

### Functional unit of Grs: molecular basis of CGA and ISQ sensing

Ligand identification of insect Grs using heterologous expression systems (e.g., human cultural cells and *Xenopus* oocytes) has often been unsuccessful and yielded little information. In the present study, we identified some ligands of silkworm Grs. The EC_50_ value of BmGr9 to D-fructose from the present study (aM level) is significantly lower than that previously reported by Sato *et al*. (mM level) (Sato et al., 2011). They did not observe BmGr9 response to D-glucose, sucrose, and *myo*-inositol up to 100 mM using calcium imaging assay in the HEK293T cells (Sato et al., 2011). It is possible that differences in Gr expression levels and calcium indicators may be responsible for the differences in susceptibility, but further investigation is needed to fully understand the cause of this discrepancy. Meanwhile, the CRISPR/Cas9-mediated KOs of BmGr6 and BmGr9 demonstrated that they could confer the MP attomolar sensitivity to the mulberry compounds (Fig. 5) in partial agreement with the results from calcium imaging using HEK293T cells (Fig. 2).

Our results confirm that single Gr can fully function as a receptor *in vitro*; nevertheless, it is evident that heterologous expression of single Gr does not fully reconstitute its function *in vivo*. For example, although BmGr6 served as a CGA receptor with attomolar sensitivity in HEK-293T cells (Fig. 2), the KO of BmGr6 did not influence the MP sensitivity of CGA (Fig. 5). This is not due to functional redundancy with other Grs, as the KO of BmGr9 resulted in a drastic decrease in CGA sensitivity (Fig. 5). It is possible that BmGr6 in MP neurons does not function as a homomeric complex but function as a heteromeric complex by coupling with other Grs, as proposed in Ors in general and in some Grs (Del Mármol et al., 2021; Jiao et al., 2008; Shim et al., 2015; Vosshall et al., 1999), resulting in different ligand spectra and sensitivity from the homomeric complex. Otherwise, the difference in the expression levels and other factors between HEK293T cells and MP neurons may affect the contribution of BmGr6 to the CGA sensitivity in MP neurons. Another example is that both BmGr6 and BmGr9 are indispensable for ISQ sensing with attomolar sensitivity (Fig. 5), suggesting a synergism between BmGr6 and BmGr9 to realize the attomolar sensitivity. In this case, too, one possibility is that the two Grs couples in a functional heteromeric complex. Altogether, our results suggest that BmGr6 and BmGr9 function as a heteromeric complex in MP neurons. Although the heterologous expression of single Grs significantly contributed to the screening for Grs with attomolar sensitivity to CGA and ISQ, its utility for dissecting *in vivo* molecular gustatory systems is limited. We cannot rule out the possibility that Grs with lower sensitivities to CGA and ISQ than BmGr6 and BmGr9 and even those with no sensitivities *in vitro* are involved in attomolar sensing in MP by synergism with the two Grs. To understand insect gustatory systems, it is required to decipher functional combinations of Grs and the mechanism determining their ligand spectra.

### Putative fructose and sugar Grs in the silkworm act as plant phenolics receptors

We demonstrate that BmGr6 and BmGr9 are responsible for ultrahigh-sensitive detection of CGA and/or ISQ. The clades to which BmGr6 and BmGr9 belong are known as sugar receptor clades (Wanner and Robertson, 2008). Nevertheless, we found that they are highly tuned to CGA and ISQ *in vitro* and responded to sugars at relatively high concentrations (Fig. 2). As there was no drastic decrease in the feeding amount of the artificial diet in BmGr6-KO and BmGr9-KO larvae in comparison with the WT (Fig. 4C), it is unlikely that BmGr6 and BmGr9 also act as sucrose receptors in the MG.

Unfortunately, we could not provide direct evidence for that because we failed to record spikes from sugar-sensing neurons in the MG of the N4 silkworm strains (both WT and mutants), probably because of their small amplitudes. Our data suggest that the two Grs act as plant phenolics receptors rather than sugar receptors in silkworm larvae’s gustatory system. Phenolics are the most widely distributed plant secondary metabolites and serve as stimulants and deterrents toward insects in feeding and oviposition (Singh et al., 2021). Some Grs in the putative sugar and fructose receptor clades may be receptors for plant phenolics which are responsible for host plant selection in herbivorous insects.

### The attomolar sensitivity is required for sensing leaf surface compounds in host plant selection

The experiments using BmGr6-KO and BmGr9-KO larvae provided further insights into the host recognition by MP. BmGr6-KO larvae lost their ability to sense ISQ but could still detect the remaining two test bite inducers (CGA and β-sitosterol) with the same ultrahigh sensitivity as the WT larvae (Fig. 5 and S3). Their low probability of test biting toward mulberry leaves (Fig. 4) confirmed the obligate authentication of the three compounds by MP. BmGr9-KO larvae have ∼1,000-fold lower sensitivity toward CGA and ISQ. However, they exhibited high sensitivity even at the femtomolar level (Fig. 5), resulting in a lack of host recognition ability with high probability (Fig. 4E). This indicates that the femtomolar sensitivity is insufficient, and the attomolar sensitivity is required to sense leaf surface secondary compounds for host recognition. We recently showed silkworm MP’s attomolar sensitivity toward feeding deterrents, such as coumarin and strychnine nitrate (Shii et al., 2021). These ultrahigh sensitivities underlie feeding decisions by herbivorous larvae at the leaf surface.

In summary, we identified the two silkworms Grs responsible for the ultrahigh sensitivity in MP to sense leaf surface host phenolic compounds and recognize host leaves. Mechanistic insight into the gustatory system in herbivorous insects, especially on a functional unit of Grs, is needed to know how their Grs have adapted their tunings to plant secondary metabolites and their contribution to the evolution of feeding preferences.

## Material and methods

### Chemicals

| chemicals | source | identifer | CAS RN |
| --- | --- | --- | --- |
| chrologenic acid | Nacalai tesque | 08416-14 | 327-97-9 |
| isoquercetrin | Extrasynthese | 1327S | 482-35-9 |
| β-sitosterol | Abcam | ab143122 | 83-46-5 |
| D-glucose | Fujifilm Wako Pure Chemical Corp. | 049-31165 | 50-99-7 |
| D-fructose | Fujifilm Wako Pure Chemical Corp. | 127-02765 | 57-48-7 |
| sucrose | Fujifilm Wako Pure Chemical Corp. | 196-00015 | 57-50-1 |
| *myo*-inositol | Fujifilm Wako Pure Chemical Corp. | 092-00282 | 87-89-8 |

### Insect rearing

The silkworm *Bombyx mori* larvae (N4 strain and Kinshu × Showa hybrid) were reared at 25°C on an artificial diet Silkmate 2M (Nihon-Nosan Co. Ltd.) under a 12 h light–12 h dark photoperiod. The N4 larvae were used for experiments in Fig. 2, 4, 5, S3, and S4. The Kinshu × Showa larvae were used for immunostaining in Fig. 3.

### Phylogenetic analysis

The lepidopteran Gr phylogeny was generated using full-length amino acid sequences of Grs from *Bombyx mori* (Guo et al., 2017), *P. xylostella* (Engsontia et al., 2014), *H. melpomene* (Briscoe et al., 2013), PxutGr1 (Ozaki et al., 2011), and PrapGr28 (Yang et al., 2021). The sequences were aligned with MAFFT v.7 (http://mafft.cbrc.jp/alignment/server/). The IQ-TREE web server (http://iqtree.cibiv.univie.ac.at/) generated the maximum-likelihood tree using the JTT+F+I+G4 model. The phylogenetic tree was visualized using iTOL (https://itol.embl.de/) (Letunic and Bork, 2021).

### cDNA cloning of *Bombyx* Grs

cDNA of BmGr6, BmGr7, BmGr9, and BmGr10 were cloned into the pcDNA3.1 vector (Thermo Fisher Scientific) as reported previously (Kikuta et al., 2016; D. Mang et al., 2016). According to the nucleotide sequences reported by Guo *et al*. (Guo et al., 2017), cDNA of BmGr4, BmGr5, and BmGr8 were codon-optimized for mammalian cell expression, artificially synthesized (gBlocks^®^, Integrated DNA Technologies), and inserted into the EcoRV site of pcDNA3.1 vector using GeneArt^®^ Seamless Cloning and Assembly Enzyme Mix (Thermo Fisher Scientific).

### Cell culture and transfection

HEK293T cells were cultured in Dulbecco’s modified Eagle medium (DMEM, Thermo fisher scientific) supplemented with 10% fetal bovine serum (BioWest), 4 mM GlutaMAX (Thermo Fisher Scientific), 100 units/ml penicillin (Meiji Seika Pharma Co., Ltd.), and 100 mg/ml streptomycin (Meiji Seika Pharma Co., Ltd.) with 5% CO_2_ at 37°C. For preparing transfection solutions, 500 µL Opti-MEM (Thermo Fisher Scientific) were vortexed with 4–6 µg polyethylenimine (PEI Max, Polysciences, Inc., Warrington, PA) and incubated for 5 min. Afterward, 2 µg of an expression vector was added, vortexed vigorously, and incubated for 20 min. Approximately 50%–70% confluent HEK293T cells were seeded on a coverslip in a six-well plate and transfected with the transfection solution for 4 h with 5% CO_2_ at 37°C. The media was replaced with DMEM, and the cells were incubated for 48 h.

### Calcium imaging

For preparing the loading solution of Ca^2+^ indicator Fluo-4 AM (Thermo Fisher Scientific), 3 µL of 1 mM Fluo-4 AM (stock solution in DMSO) and 30 µL of PowerLoad Concentrate (100X, Thermo Fisher Scientific) were vortexed vigorously. The mixture (5 µL) was added into a six-well plate containing 500 µL Hank’s buffered saline solution (HBSS, 137 mM NaCl, 5.4 mM KCl, 0.3 mM Na_2_HPO_4_, 0.4 mM KH_2_PO_4_, 4.2 mM NaHCO_3_, 1.3 mM CaCl_2_, 0.5 mM MgCl_2_, and 0.4 mM MgSO_4_, pH 7.4) incubated for 10 min at 37°C (final conc. ∼1 µM Fluo4-AM and ∼1X PowerLoad). After loading, the solution was replaced with HBSS containing 0.75 mM Probenecid (Thermo Fisher Scientific). After 1 h incubation, the cells were used for calcium imaging experiments.

Calcium imaging was performed with a flow system under the fluorescence microscope BX53 (Olympus) equipped with a MetaView imaging system (Molecular Devices). A coverslip on which transfected cells were seeded was inverted and put on a self-manufactured chamber with a 5-mm-wide flow channel. The HBSS buffer was superfused at a proper flow rate in a silicon tube with peristaltic pumps (PERISTA pump, ATTO). 200 µL of ligand solution was loaded into the tube with small bubbles back and forth to separate it from HBSS. CGA (Nacalai tesque, Kyoto, Japan), ISQ (Extrasynthese, Lyon, France), D-glucose, D-fructose, sucrose, and *myo*-inositol (Fujifilm Wako Pure Chemical Corp., Osaka, Japan) were dissolved and serially-diluted in HBSS at time of use. β-sitosterol (Abcam, Cambridge, UK) was dissolved in a 45% solution of 2-hydroxypropyl-β-cyclodextrin at 37°C to prepare stable aqueous solutions. The time-lapsed fluorescence images (1 Hz) were processed using Fiji (Schindelin et al., 2012), and the fluorescent ratio (ΔF/F) was calculated as peak intensity divided by basal intensity (mean of 10 flames before ligand perfusion). The EC_50_ value was calculated by fitting using the four-parameter logistic model (Prism ver.8, Graph-Pad).

### Immunostaining

The heads of the fifth instar larvae (Kinshu × Showa hybrid) were fixed with 4% paraformaldehyde/phosphate-buffered saline (PBS, 137 mM NaCl, 2.7 mM KCl, 10 mM Na_2_HPO_4_, 1.8 mM KH_2_PO_4_, pH 7.4) overnight at 4°C. The fixed heads were dissected, and the maxilla was collected and immersed into a cryoprotection solution (10% sucrose/PBS at room temperature (RT) for 3 h; 20% sucrose/PBS at 4°C overnight; 30% sucrose/PBS at 4°C overnight), embedded in Tissue-Tek O.C.T. Compound (Sakura Finetek Japan Co., Ltd) and stored at −80°C. The embedded specimens were sectioned at 14 µm thickness via cryostat (CM3050S, Leica, Wetzlar, Germany), mounted on the coated slide glasses (MAS-03, Matsunami Glass, Osaka, Japan), and air dried at RT for a day. After 5 min washes with Tris-NaCl-Tween buffer (TNT; 0.15 M Tris-HCl, 0.01 M NaCl, 0.05% Tween20), the sections were permeabilized with 0.1% TritonX-100/TNT for 5–10 min, blocked with 2% bovine serum albumin (BSA)/TNT for 30 min, and incubated with primary antiserum raised elsewhere (Mang et al., 2016; D. Mang et al., 2016) (anti-BmGr6 and anti-BmGr9 antisera in 2% BSA/TNT at a 1:100 dilution) for 12 h at 4°C. The specificities of these antisera were confirmed by testing cross-reactivity of closely related BmGr proteins (Mang et al., 2016; D. Mang et al., 2016). After three 5 min washes with TNT, the sections were incubated in a secondary antiserum (an antimouse IgG conjugated with Alexa Fluor 488, Molecular Probes, USA), at a 1:1,000 dilution in TNT for 12 h at 4°C. After three TNT washes for 5 min, the sections were counterstained with 4, 6-diamidino-2-phenylindole (l mg/mL, Sigma Aldrich, Schnelldorf, Germany) for 5 min at RT. After 5 min of TNT wash, the sections were mounted in 90% glycerol/PBS containing 1, 4-diazabicyclo[2.2.2] octane at RT and observed under fluorescence microscopy (BX53, Olympus, Tokyo, Japan).

### Generation of Gr-KO silkworm strains using CRISPR/Cas9 genome editing

We used the Alt-R® CRISPR-Cas9 system (Integrated DNA Technologies) to generate silkworm strains carrying null mutations in *BmGr6* or *BmGr9* genes. CRISPRdirect (Naito et al., 2015) identified unique target sequences for guide RNAs using *Bombyx mori* genome assembly as a reference. The off-target search against silkworm genomic sequence confirmed that the designed sgRNAs were highly specific to the target sequences and had only one match (no predicted off-target sites) in 20mer+PAM and 12mer+PAM search. Therefore, we consider that do not cause off-target effects theoretically. Alt-R® crRNAs (for BmGr6: 5′-GTATTCGGCGGCGCTGGGAA<u>TGG</u>-3′; for BmGr9, 5′- CTGGTGGCGGAGATAGGAGT<u>GGG</u>-3′), tracrRNA, and S.p. Cas9 Nuclease V3 were purchased from Integrated DNA Technologies. CRISPR-Cas9 ribonucleoprotein (RNP) complexes containing 10 µM crRNA:tracrRNA duplex and 300 ng/µL Cas9 enzyme were prepared according to a user-submitted protocol entitled “CRISPR-Cas9 RNP delivery—*B. tryoni* microinjection—A Choo” on the Integrated DNA Technologies website. Newly laid eggs of the WT N4 strain were aligned and glued onto a sliding glass, and RNP complexes were injected using IM 300 Microinjector (NARISHIGE) within 5–8 h after oviposition. Generation 0 (G_0_) moths were crossed with WT moths. To identify Generation 1 (G_1_) broods carrying a heterozygous Cas9-induced mutation, genomic DNA extracted from ∼10 G_1_ eggs were used for the T7 Endonuclease I (T7EI) assay with Alt-R^®^ Genome Editing Detection Kit (Integrated DNA Technologies). T7EI assay-positive G_1_ moths were subjected to sibling mating or crossed with WT moths. Mutations were identified by DNA sequencing of genomic PCR (gPCR) products. Mutant strains carrying a homozygous null mutation were generated by intercrossing G_2_ or G_3_ moths.

### Feeding behavior assays

The feeding behavior assays were performed as described previously (Tsuneto et al., 2020). For feeding assay, three to five fifth instar larvae (Day 0) were put on individual Petri dishes, and the larval mass before and after 1.5 h feeding was measured. An increase in larval mass was regarded as the amount of food intake because no feces were observed after feeding. We adopted an increase in body mass (%, w/w) as the index of food intake because a significant correlation between the amount of food intake and the original body mass was observed via Pearson’s correlation analysis (Fig. S4). Mulberry leaves were collected around the campus of the Tokyo University of Agriculture and Technology (Koganei, Tokyo, Japan) and stored at 4°C. To observe test bites, videos of silkworm feeding on leaves were captured by a D5100 camera (Nikon, Tokyo, Japan) equipped with a telephoto lens AF-S DX Micro Nikkor 85 mm (Nikon, Tokyo, Japan). The larvae were used 24–48 h after molting to the fifth instar. To prevent nonspecific biting due to starvation, all larvae were fed on artificial diets for 2 min before use.

### Tip recording

Tip recording was performed as described previously (Tsuneto et al., 2020). In brief, the heads of the fifth instar larvae (Day 0) were excised and used for further experiments within ∼20 min. The whole sensilla of MP were capped with a glass capillary (Erma Inc., Tokyo, Japan) filled with a tastant solution dissolved in 5 mM NaCl using a micromanipulator (NARISHIGE, Tokyo, Japan). The interval of stimulation was >1 min. When we observe desensitization in high concentration, we do not include the data and test other larvae. We wash the silver wire to remove extraneous matter especially after using high-concentrated solution. To confirm the silver wire is clean, we always test response to 5 mM NaCl first before testing stimulants. We prepared taste solutions and their serial dilution at time of use from powders (CGA and ISQ) and β-sitosterol stock solution (stored at − 20 °C). Electrical signals were amplified by a TastePROBE amplifier (Syntech, Hilversum, The Netherlands), recorded via PowerLab 2/25 (AD Instruments, Sydney, Australia), and analyzed by the LabChart software (AD Instruments). Spikes were counted from 0.1 to 1.1 s after the onset of stimulation using custom codes written in MATLAB (Mathworks). Thresholds for counting spikes were set from 0.3 to 1 mV depending on the baseline amplitude.

## Acknowledgments

The authors thank Shingo Kikuta (Ibaraki University) and Masataka G. Suzuki (The University of Tokyo) for technical assistance in calcium imaging at an early stage of this study and for establishing the CRISPR-Cas9 system. This research was supported by the Japan Society for the Promotion of Science (JSPS) KAKENHI (Grant Number 17K19261 for RS; 18J00733 for HE).

## Author contributions

H.E., K.T., and R.S. designed the research; H.E. and K.T. performed calcium imaging and electrophysiological recording studies; K.T. performed immunostaining and behavioral experiments; H.E., T.Y., and D.M. generated knockout mutants under the supervision of K.I. and S.N. H.E. wrote the manuscript with intellectual inputs from all authors.

---

**Fig. S1.**
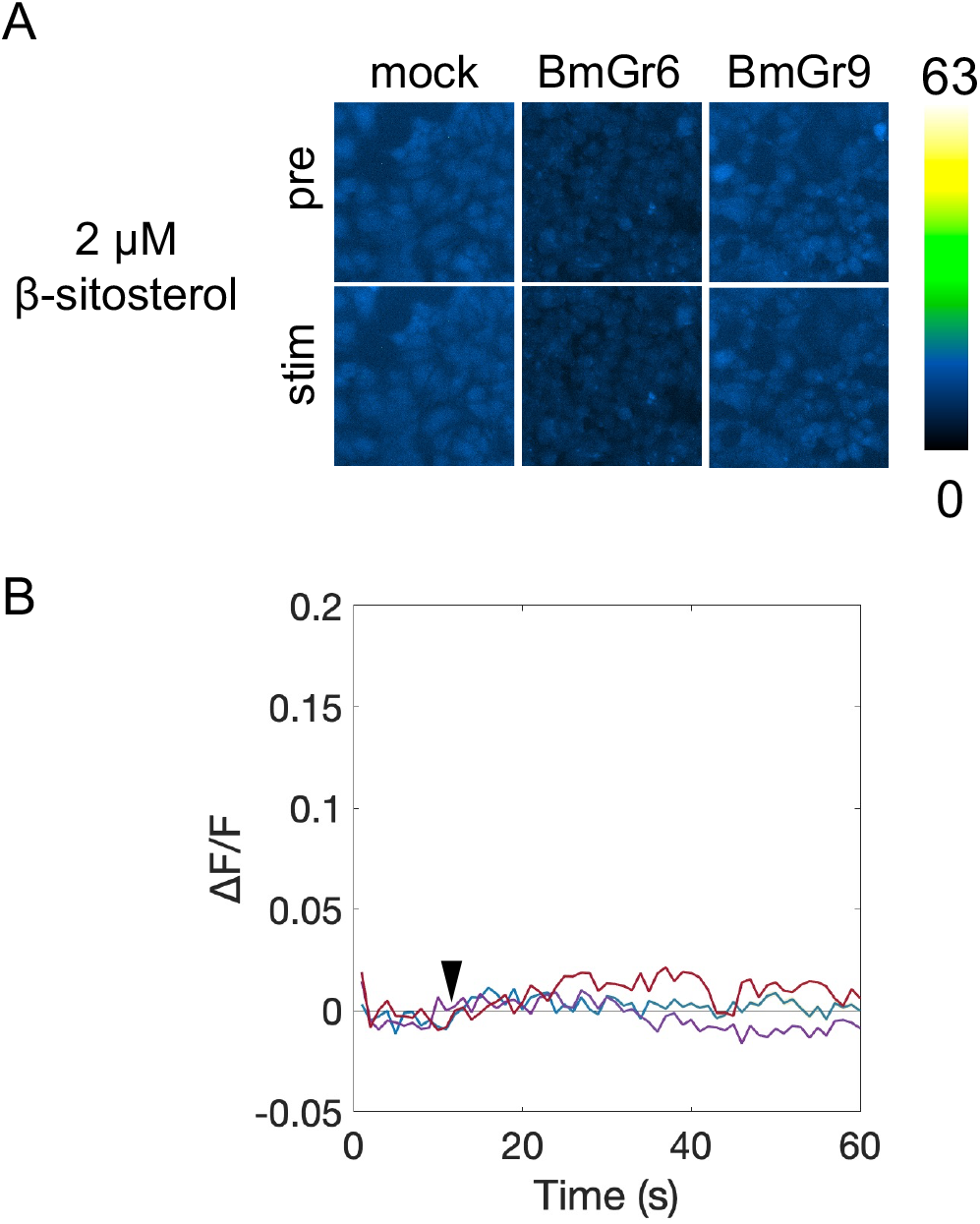
Response of BmGr6-and BmGr9-expressing cells toward β-sitosterol. (A) The representative cell images before (pre) and after (stim) stimulation by 2 µM β-sitosterol. Fluorescent intensity is color-coded according to the scale bar. (B) Time course of responses (ΔF/F) toward 2 µM β-sitosterol (n = 30). The arrowhead indicates the timing of the ligand application.

**Fig. S2.**
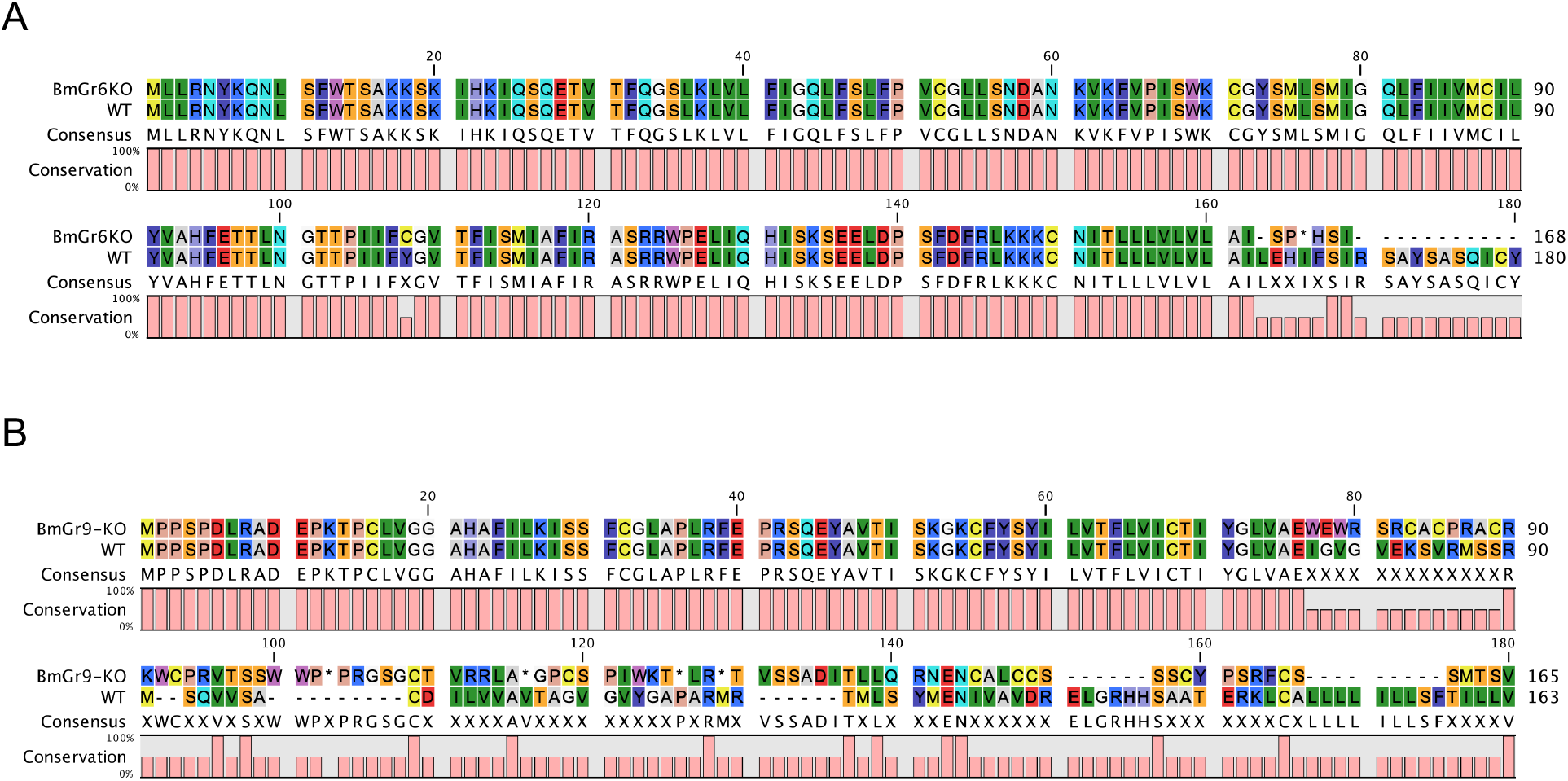
Deduced amino acid sequences of truncated BmGr6 (A) and BmGr9 (B) in knockout larvae. The sequences were aligned using CLC Sequence Viewer 8 (CLC Bio, Aarhus, Denmark).

**Fig. S3.**
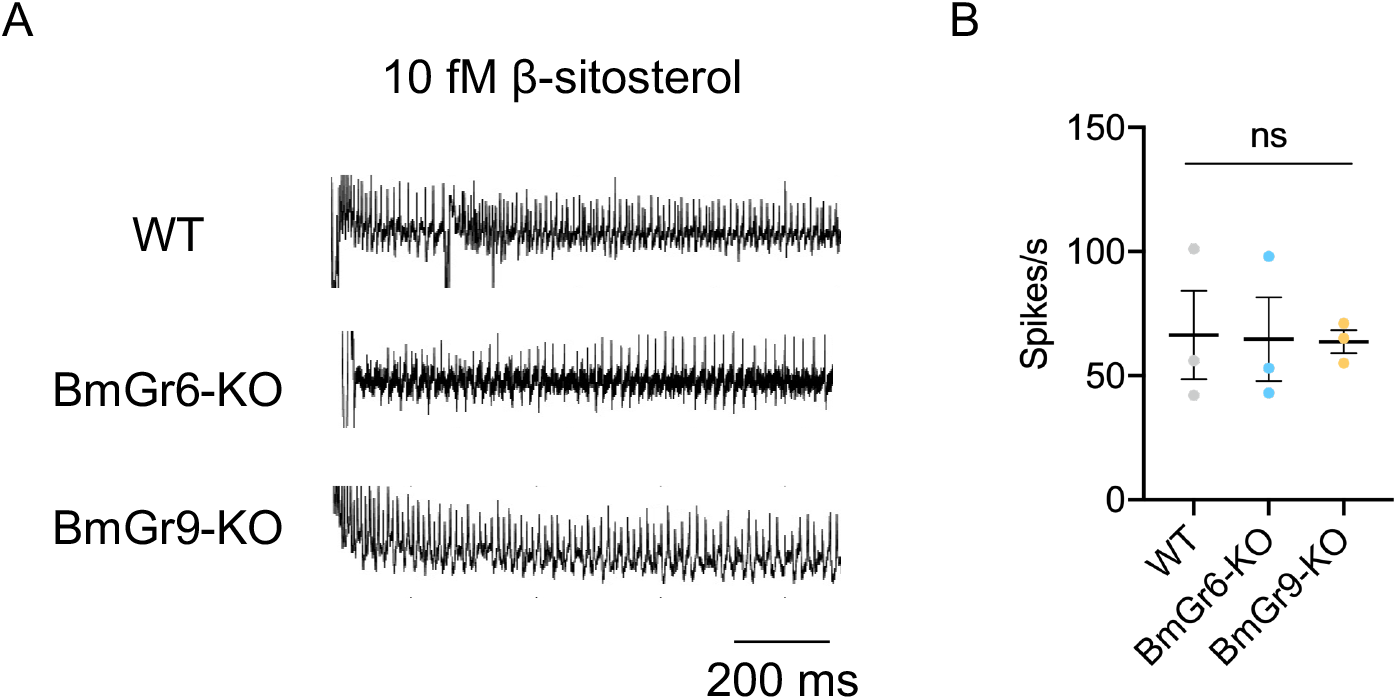
Response of the maxillary palp (MP) in BmGr6-KO and BmGr9-KO larvae toward β-sitosterol. The representative traces (A) and frequencies of action potentials from the whole MP sensilla in response to 10 fM β-sitosterol (B). Spikes were counted from 0.1 to 1.1 s after the onset of stimulation. Data are represented as the mean ± SEM (n = 3). Statistical analysis was performed via one-way ANOVA followed by the Tukey post hoc test. “ns” indicates no significant difference.

**Fig. S4.**
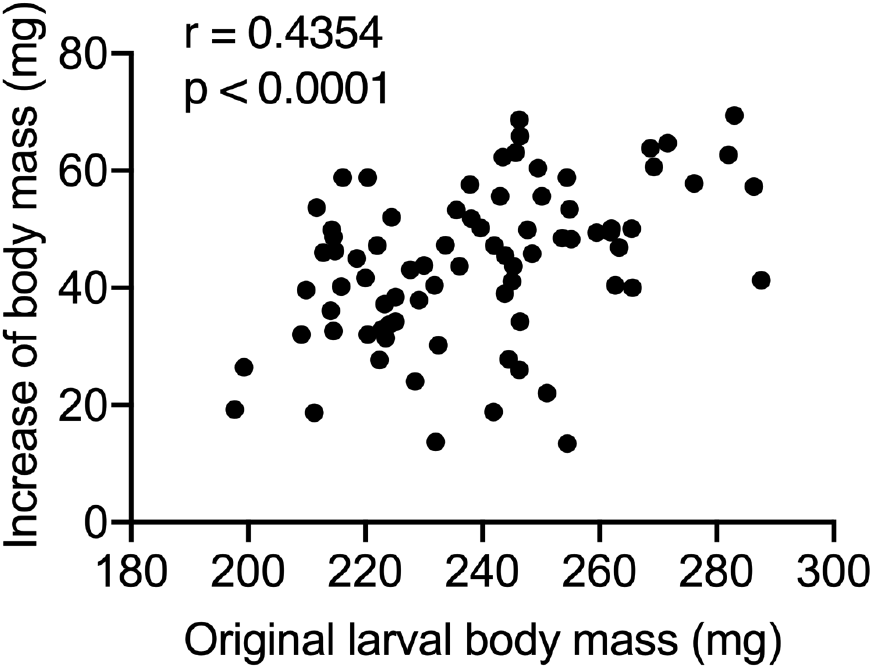
Correlation between original body mass and feeding amount. Correlation coefficient (r) using Pearson correlation analysis (n = 80).

**Table S1.**
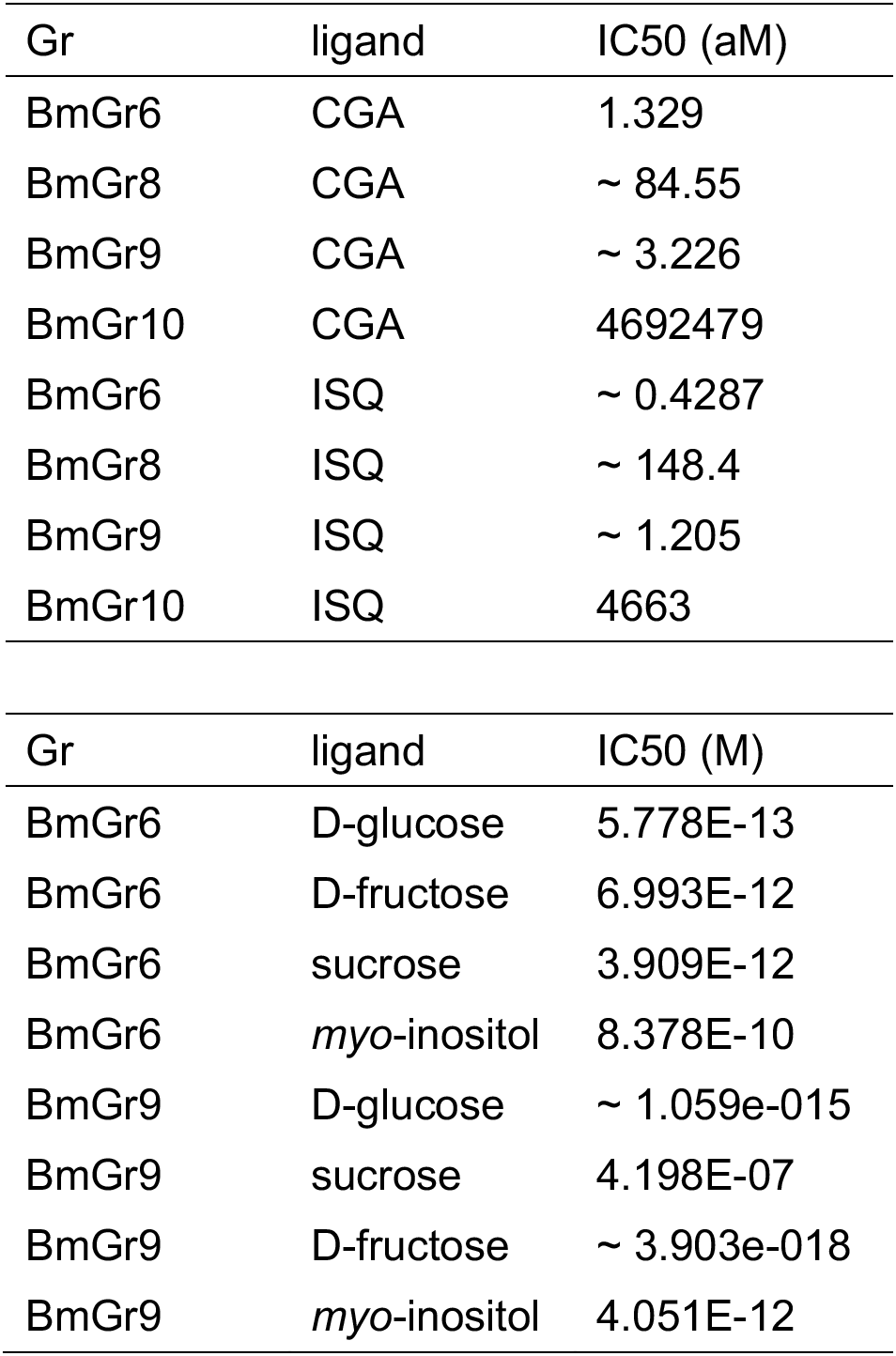
EC_50_ values of BmGrs in response to plant secondary metabolites (chlorogenic acid (CGA) and isoquercitrin (ISQ)) and sugars.

| Gr | ligand | IC50 (aM) |
| --- | --- | --- |
| BmGr6 | CGA | 1.329 |
| BmGr8 | CGA | ~ 84.55 |
| BmGr9 | CGA | ~ 3.226 |
| BmGr10 | CGA | 4692479 |
| BmGr6 | ISQ | ~ 0.4287 |
| BmGr8 | ISQ | ~ 148.4 |
| BmGr9 | ISQ | ~ 1.205 |
| BmGr10 | ISQ | 4663 |

| Gr | ligand | IC50 (M) |
| --- | --- | --- |
| BmGr6 | D-glucose | 5.778E-13 |
| BmGr6 | D-fructose | 6.993E-12 |
| BmGr6 | sucrose | 3.909E-12 |
| BmGr6 | *myo*-inositol | 8.378E-10 |
| BmGr9 | D-glucose | ~ 1.059e-015 |
| BmGr9 | sucrose | 4.198E-07 |
| BmGr9 | D-fructose | ~ 3.903e-018 |
| BmGr9 | *myo*-inositol | 4.051E-12 |

